# A Multidimensional Framework for Behavioral Persistence: Dissociable Dimensions of Effort, Endurance, and Sequence Stability in Mice

**DOI:** 10.64898/2026.03.02.709169

**Authors:** Tianyu Cao, William R. Johnston, Stephen Christensen, Samuel Crouse, Qian-Quan Sun

## Abstract

Behavioral persistence, the maintenance of goal-directed action despite obstacles, is a fundamental adaptive process, yet its scientific study remains fragmented across disciplines with disparate operational definitions. Here, we introduce the Persistence Spectrum (PERCS) framework, a five-dimensional model deconstructing persistence into Perseverance of Effort (P), Strategic Endurance (E), Resistance to Extinction (R), Temporal Consistency (C), and Repetitive Sequence Stability (S). Using programmable operant schedules via the FED3 system, we induced a continuum of persistent food-seeking in mice across four paradigms: Fixed Ratio (FR), Alternating 2×2 and 5×5, and Random Progressive Ratio (RPR). To objectively identify persistence periods, we developed a session-specific Gaussian mixture model, providing a data-driven alternative to arbitrary frequency thresholds. We found that persistence bouts were fundamentally driven by unrewarded effort, not reward delivery: instantaneous frequency for incorrect pokes significantly exceeded that for correct pokes in FR, 2×2, and 5×5 (all p < 0.05). This pattern was most pronounced in the high-demand RPR schedule, where high-rate poking continued unabated even after successes, directly challenging reinforcement-centric models and providing strong empirical support for frustration theory. Linear mixed-effects modeling revealed that frequency increased with consecutive unrewarded pokes across all paradigms, with a quadratic (inverted-U) relationship specific to FR and 2×2 (p < 0.001), suggesting effort invigoration followed by strategic disengagement in simpler tasks, whereas high-demand schedules maintained linear increases. Notably, the collapse of Sequence Stability (S) in RPR is partly constrained by the task’s environmental contingency: random reward rules mathematically restrict stable sequence learning. Aggregate analysis showed total poke counts and unrewarded effort scaled with task difficulty (all p < 2e-16), yet pellet retrieval rates remained stable, indicating goal achievement despite increased challenge. Crucially, PERCS profiles were robust across independent and continuous training histories, demonstrating they reflect stable phenotypes shaped by current contingencies rather than training artifacts. Application of PERCS revealed distinct fingerprints: FR produced near-zero P, E, R, and C but maximal S, characteristic of an efficient habit; 2×2 and 5×5 elevated P, E, and R while reducing S; and RPR generated highest P and R, lowest S, and marked inter-individual variability. These findings demonstrate that operant schedules dissociably shape distinct persistence dimensions, with unrewarded effort acting as a key motivational trigger, positioning frustrative nonreward as a primary engine of persistent behavior. The PERCS framework provides a unified, quantitative language for characterizing persistent behavior across species and paradigms, offering a powerful tool for linking dimensions to neural circuits and understanding their breakdown in neuropsychiatric disorders.

## Introduction

Behavioral persistence, the maintenance of goal-directed actions despite obstacles, delay, or diminishing returns, is a fundamental adaptive process across species, essential for survival, foraging, and long-term achievement. Despite its centrality, the scientific study of persistence is conceptually fragmented, scattered across disciplines with disparate operational definitions and terminologies that hinder a unified, mechanistic understanding (Nevin and Grace 2000); (Amsel and Roussel 1952). In comparative psychology, persistence is often studied through specific, narrow lenses. Behavioral momentum theory conceptualizes it as resistance to change when reinforcement is disrupted or withheld (Nevin and Grace 2005, Podlesnik and Shahan 2008).

Progressive ratio (PR) schedules reduce it to a single motivational “breakpoint” (Hodos 1961); (Richardson and Roberts 1996, Bradshaw and Killeen 2012), while foraging models describe it as the scalable continuation of feeding under threat. In parallel, human psychology has developed constructs like grit, defined as perseverance for long-term goals (Duckworth, Peterson et al. 2007), and delay of gratification (Mischel, Shoda et al. 1989), which focus on trait-level or cognitive self-regulation but lack direct neurobiological translatability. Clinical research further complicates the picture by studying pathological extremes, such as compulsivity in obsessive-compulsive disorder (Graybiel and Rauch 2000) or amotivation in depression (Treadway and Zald 2011), and cognitive rigidity in autism spectrum disorder (Geurts, Corbett et al. 2009, Lord, Brugha et al. 2020),without a dimensional framework that connects these states to adaptive persistence. This fragmentation presents a fundamental challenge: we lack a common operational vocabulary and a controlled experimental paradigm capable of inducing and dissecting the full continuum of persistent states, from efficient goal-pursuit to maladaptive perseveration(Eisenberger 1992, Carver and Scheier 1998).

Recent technological advances in behavioral monitoring offer a path forward. The Feeding Experimentation Device version 3 (FED3) has emerged as a transformative tool for high-resolution, home-cage measurement of feeding microstructure and operant behavior, minimizing stress and enabling rich, longitudinal phenotyping (Matikainen-Ankney, Earnest et al. 2021). It has been used descriptively to study threat-suppressed feeding (de Araujo Salgado, Li et al. 2023), effort-based motivation in obesity models (Barrett, Raineri Tapies et al. 2020), and neural correlates of consumption . However, its potential extends beyond description. Crucially, the FED3 is programmable. We propose to leverage this feature not merely to measure persistence, but to actively engineer it by systematically manipulating reinforcement contingencies(Skinner 1988) (Thorndike 1933). By designing operant schedules that parametrically vary effort cost, reward uncertainty, and sequential complexity, we can directly induce a broad spectrum of persistent behavioral states within the same subjects. This creates a controlled experimental “persistence continuum,” a necessary substrate for rigorous, multidimensional analysis.

Quantifying this continuum requires moving beyond single metrics like breakpoints or total responses (Stevens 1946; Cronbach and Meehl 1955). A mouse pressing a lever hundreds of times on a PR schedule, a bird vigilantly foraging under predation risk, and a human working diligently on a long-term project may all be described as “persistent,” but they engage qualitatively different behavioral strategies (Tinbergen 1963, 1972; McFarland and Sibly 1975; Schleidt 1974). Deconstructing persistence into its core, dissociable components is therefore essential for linking specific behavioral patterns to their underlying neural circuits and for understanding individual differences in adaptive and maladaptive behavior (Eisenberger 1992, Sayar-Atasoy, Yavuz et al. 2024)(Sayar-Atasoy et al., 2024). To address this, we introduce the Persistence Spectrum (PERCS) framework, a five-dimensional model designed to provide a unified, quantitative characterization of persistent behavior. PERCS deconstructs persistence into the core dimensions of Perseverance of Effort (P), Strategic Endurance (E), Resistance to Extinction (R), Temporal Consistency (C), and Repetitive Sequence Stability (S). Temporal Consistency (C) draws on extensive work in biological rhythms and circadian behavior (Boulos and Terman 1980, Frank, Kupfer et al. 2005, Wang and Sun 2024)(Boulos and Terman 1980; Wang and Sun 2023; Frank, Kupfer et al. 2005), while Repetitive Sequence Stability (S) builds on ethological analyses of action syntax(Markowitz, Gillis et al. 2023) (Jin, Tecuapetla et al. 2014, Wiltschko, Johnson et al. 2015, Weinreb, Pearl et al. 2024).

Our primary objective was to test the hypothesis that distinct operant schedules would differentially shape these PERCS dimensions, generating unique behavioral “fingerprints” or profiles. We predicted that schedules imposing higher effort costs (e.g., progressive ratio) would elevate P (Perseverance of Effort); schedules with unpredictable rewards or requiring complex sequences would modulate E (Strategic Endurance) and S (Sequence Stability); and the R (Resistance to Extinction) and C (Temporal Consistency) dimensions would capture fundamental differences between habit-like, automated responding and flexible, goal-directed persistence (Squire and Zola 1996, Graybiel 2008). This work establishes a new methodological paradigm: using a programmable device (FED3) to induce a persistence continuum and applying a multidimensional computational framework (PERCS) to deconstruct it. By bridging controlled operant paradigms, naturalistic ethology, and clinical concepts, this approach provides a common quantitative language for persistence research. Here, we demonstrate that a series of FED3 operant schedules, Fixed Ratio (FR), Alternating (2×2, 5×5), and Random Progressive Ratio (RPR), reliably produce a wide range of persistent feeding behaviors in mice. Our multidimensional analysis reveals that these schedules produce distinct, quantifiable PERCS profiles. For instance, the low-cost FR schedule produces high S but low P, characteristic of an efficient habit (Squire and Zola 1996, Graybiel 2008), while the high-cost RPR schedule elevates P and R but collapses S, indicative of effortful but less structured responding. These findings validate the PERCS framework as a powerful tool for mapping the adaptive landscape of persistent behavior and set the stage for future research to identify the specific neurobiological substrates governing each persistence dimension.

## Results

### Threshold Determination Using Gaussian Mixture Models

To determine the high and low instantaneous frequency thresholds used to define persistence periods objectively, we fitted a two-component Gaussian mixture model (GMM) separately for each mouse and for each training session, using the flexmix R package. This session-specific, mouse-specific approach avoids pooling data across sessions or individuals and accounts for both inter-individual and within-animal variability in poking behavior. This data-driven method provides an objective alternative to arbitrary frequency thresholds, thereby enhancing the reproducibility of “persistence” as a behavioral construct. Let *yᵢ* denote the instantaneous frequency of the ith poke within a given mouse and session. We assumed that *yᵢ* arises from a mixture of two normal distributions, *yᵢ ∼ π₁N(μ₁, σ₁²) + π₂N(μ₂, σ₂²)*, where *π₁* and π₂ = 1-π₁ are the mixture proportions, and *μₖ* and *σₖ* denote the mean and variance of component k, for k = 1,2 . Although the true distribution of instantaneous frequencies is unknown, this two-component Gaussian mixture model provided a reasonable and robust separation between low-and high-frequency pokes. Model parameters (*πₖ, μₖ, σₖ²*) were estimated using the expectation–maximization (EM) algorithm. For each poke i, the posterior probability of belonging to component k was computed as *Pr(zᵢ = k | yᵢ) = (πₖ N(yᵢ | μₖ, σₖ²)) / (∑ⱼ πⱼ N(yᵢ | μⱼ, σⱼ²))*, where *zᵢ* is a latent component indicator. Each poke was assigned to the component with the higher posterior probability. The high instantaneous frequency threshold was defined as High bound = (max{yᵢ | zᵢ = low}1 + min{yᵢ | zᵢ = high}) / 2, and the low instantaneous frequency threshold was defined as 𝐿*ow* 𝑏*ound* = 𝜇*_low_*. These session-specific thresholds were then used to identify persistence periods. To visualize the mixture model and the resulting thresholds, we added fitted Gaussian envelopes to the histograms of instantaneous frequencies. An example from a single session of a randomly selected mouse from the 2×2 paradigm shows that the two-component Gaussian mixture model, while not perfectly capturing all features of the data, provides sufficiently distinct components to determine the threshold for persistence periods (Supplementary Figure 1).

### Expression of Persistence Behavior Across Paradigms

To dissect the architecture of persistent pursuit, we aligned poking behavior to the onset of persistence and non-persistence periods across schedules of escalating cost (Figure 1B). In the low-effort FR schedule, poking was characterized by sparse, low-frequency events. Conversely, the high-cost RPR schedule evoked the highest overall poking density, marked by sustained high instantaneous frequencies, indicating maximal energetic investment. Relative to the intermediate 2×2 and 5×5 schedules, RPR mice initiated persistence periods more frequently, though each bout was shorter in duration. Across all paradigms, persistence periods were characterized by a striking elevation in poke frequency compared to equivalent non-persistence epochs. Critically, high-frequency poking clusters were composed almost exclusively of incorrect pokes, revealing that these bouts represent phasic bursts of unrewarded, cost-tolerant effort rather than reward-oriented correct responses.

**Figure 1:**
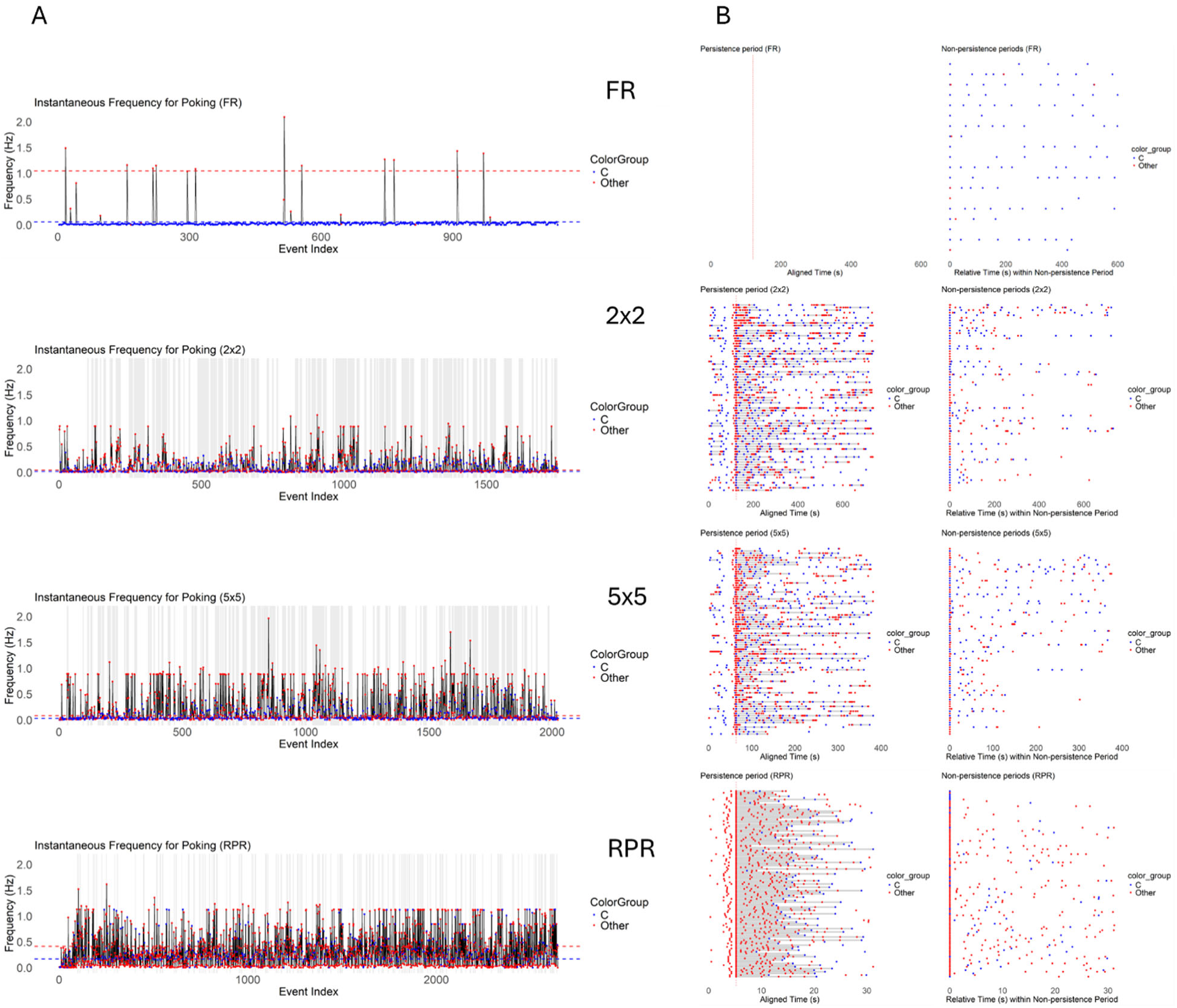
Instantaneous poke frequency and raster dynamics across feeding paradigms. The figure consists of four rows and two columns. Each row corresponds to one feeding paradigm (from top to bottom: FR, 2×2, 5×5, and RPR). For each paradigm, **A (left column)** and **B (right column)** present complementary views of poke behavior from the same representative mouse. **(A) Instantaneous frequency of poking.** For each paradigm, the instantaneous poke frequency was calculated as the reciprocal of the inter-poke interval (time between the current poke and the preceding poke). Each dot represents a single poke plotted over time. Blue dots indicate correct pokes, and red dots indicate other pokes. Gray shaded regions denote persistence periods (persistence bouts). **(B) Raster-like plot aligned to behavioral state transitions.** Raster plots show poke timing across persistence and non-persistence periods. Each dot represents a single poke, with time on the x-axis. For persistence periods, onset times were aligned to the vertical dashed line. For non-persistence periods, the beginning of each non-persistence period was aligned to zero. This alignment allows direct comparison of poke dynamics relative to the onset of each behavioral state. Blue and red dots again indicate correct and other pokes, respectively, and gray shading denotes persistence bouts.

### Persistence Periods Are Driven by Unrewarded Effort

To determine whether persistence periods represent bursts of unrewarded effort, we compared instantaneous poking frequencies for correct versus incorrect pokes within each schedule (Figure 2A). In the FR, 2×2, and 5×5 schedules, instantaneous frequency was significantly lower for correct pokes than for other pokes (FR: t₍₄₎ = −9.76, p = 0.0006, mean difference = −0.705; 2×2: t₍₄₎ = −21.98, p < 0.0001, mean difference = −0.108; 5×5: t₍₄₎ = −3.50, p = 0.025, mean difference = −0.148), confirming that high-frequency clusters are composed primarily of unrewarded responses. Notably, this pattern held even in the FR schedule, which lacks formal persistence periods, suggesting that the drive for high-frequency poking is not contingent on schedule complexity. In sharp contrast, the RPR schedule revealed no significant difference between correct and other pokes (t₍₄₎ = 2.49, p = 0.067, mean difference = 0.034), with incorrect pokes exhibiting a marginally lower mean frequency than correct pokes. This divergence reflects the unique contingency of the RPR schedule, in which reward delivery depends on a cumulative poke count across both ports, rather than on consecutive correct responses.

**Figure 2:**
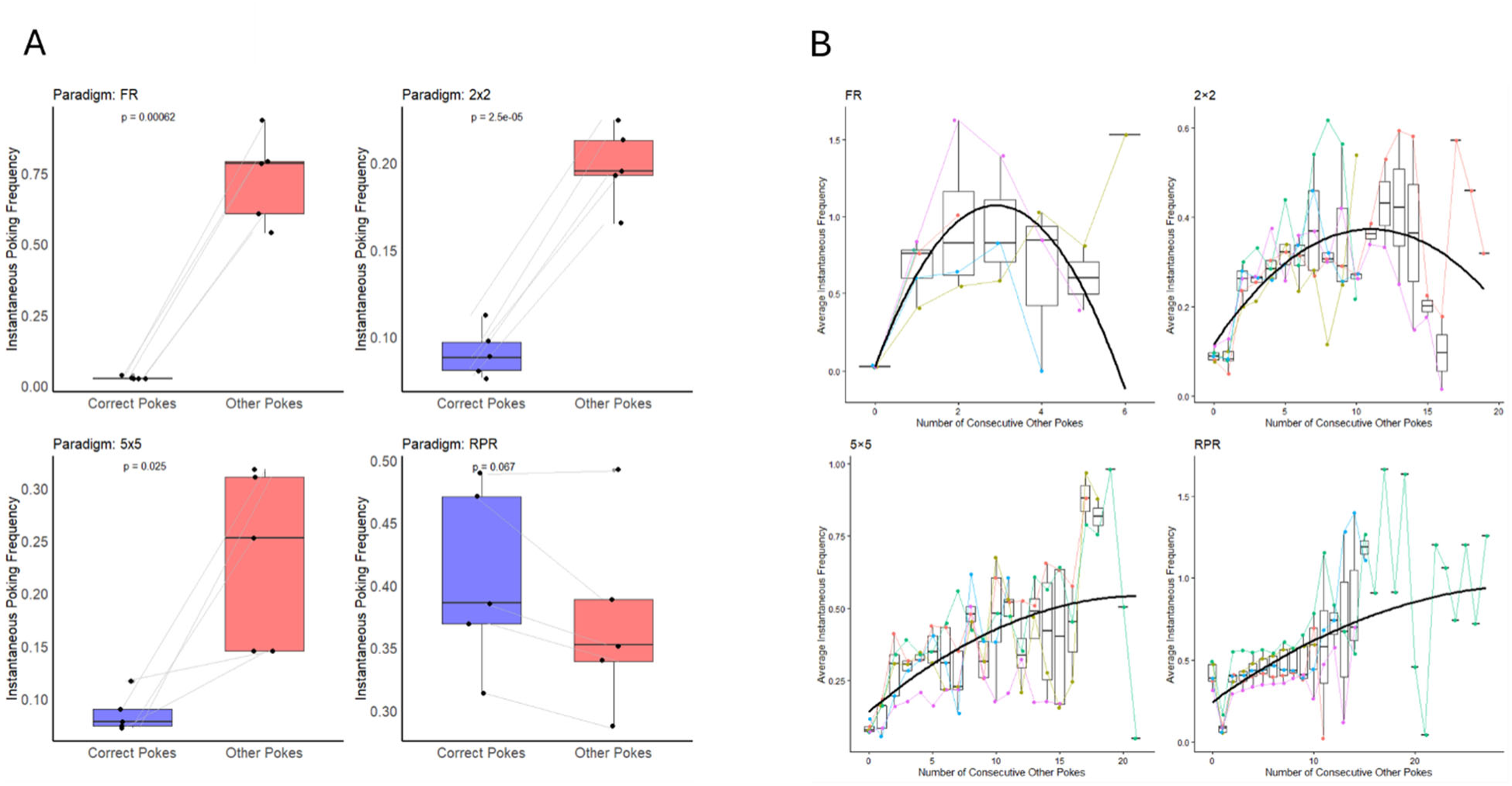
Instantaneous frequency differences between poke types and modulation by consecutive unrewarded pokes across paradigms. The figure contains two main panels: **(A, left)** and **(B, right)**, showing analyses across the four feeding paradigms (FR, 2×2, 5×5, and RPR). **(A) Paired comparison of instantaneous frequency for correct and other pokes.** For each mouse within each paradigm, the average instantaneous frequency was calculated separately for correct and other (unrewarded) pokes. Each dot represents one mouse, and gray lines connect paired values from the same mouse. Statistical significance was assessed using a two-sided paired t-test comparing correct versus other pokes. **(B) Instantaneous frequency as a function of consecutive other pokes.** For each mouse within each paradigm, the average instantaneous frequency is plotted against the number of consecutive other (unrewarded) pokes. Dots represent the mean instantaneous frequency at each consecutive other poke value, and lines connect data points from the same mouse. The black solid curve shows the quadratic fit from the mixed-effects model, reflecting the overall trend across mice.

Consequently, rewards do not reliably terminate a persistence bout, and mice continue poking at high rates even after successes, accounting for the frequent but brief persistence periods observed in Figure 1. Together, these results are consistent with frustration theory(Amsel and Roussel 1952), wherein the omission of an expected reward energizes ongoing behavior, driving phasic bursts of unrewarded effort. This finding directly challenges reinforcement-centric models of persistence, which would predict that rewarded actions are the primary drivers of behavior. Instead, our data position frustrative nonreward as a primary engine of persistent responding.

### Influence of Consecutive Unrewarded Pokes on Effort

To further examine how unrewarded actions influence poking behavior, we introduced a new variable, “Number of Consecutive Other Pokes,” which counts how many consecutive pokes do not yield a pellet. Specifically, this variable is reset to zero whenever a correct poke resulting in a pellet occurs or when a pellet is retrieved. Counting begins only after the first unrewarded (“other”) poke following a correct poke or pellet retrieval, and it increments with each subsequent consecutive unrewarded poke until a rewarded action occurs. This measure allows us to investigate how mice adjust their effort when repeated pokes fail to produce a reward (Figure 2B). We fitted a linear mixed-effects model separately for each paradigm: *Yᵢⱼ = β₀ + β₁Xᵢⱼ + β₂Xᵢⱼ² + bᵢ + ɛᵢⱼ*, where *Yᵢⱼ* = average instantaneous frequency for mouse i at a given consecutive-other-poke count, *Xᵢⱼ* = number of consecutive other pokes, *β₀, β₁, β₂* = fixed effects (intercept, linear, quadratic slope), *bᵢ* = random intercept for mouse i, and *ɛᵢⱼ* = random error. To account for heteroscedasticity, residual variances were allowed to differ across levels of *Xᵢⱼ* using a *varIdent* variance structure. Specifically,

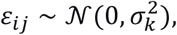

where the residual variance *σ*^2^*_k_* depends on the level 𝑘of *Xᵢⱼ*. This heteroscedastic variance structure was implemented for all paradigms.

In Table 1, we can see that for all four paradigms, the slope of the linear term (consec_Other) was positive and statistically significant (p < 0.05), indicating effort increases as failures rise. However, the quadratic (inverted-U) term was only significant in the FR and 2×2 paradigms (p < 0.001). We found a quadratic (inverted-U) relationship in simpler tasks (FR, 2×2), revealing an initial effort invigoration followed by strategic disengagement (p < 0.001), whereas high-demand tasks (5×5, RPR) maintained linear effort increases, likely driven by the energizing effects of frustrative nonreward. Overall, these results imply that for most paradigms, average instantaneous poking frequency increases approximately linearly with consecutive other poke counts, while in some cases (e.g., FR, 2×2) the relationship exhibits a nonlinear, inverted-U shape.

**Table 1:** Fixed Effects from Linear Mixed-Effects Models Across Paradigms.

| Paradigm | Fixed Effect | Estimate | Std. Error | DF | t-value | p-value |
| --- | --- | --- | --- | --- | --- | --- |
| FR | Intercept | 0.02742 | 0.00224 | 16 | 12.239 | <0.001 |
|  | consec_Other | 0.72071 | 0.08097 | 16 | 8.901 | <0.001 |
|  | (consec_Other)^2 | -0.12415 | 0.01978 | 16 | -6.278 | <0.001 |
| 2×2 | Intercept | 0.11582 | 0.03602 | 63 | 3.216 | 0.0021 |
|  | consec_Other | 0.04661 | 0.00894 | 63 | 5.212 | <0.001 |
|  | (consec_Other)^2 | -0.00212 | 0.00052 | 63 | -4.061 | 0.0001 |
| 5×5 | Intercept | 0.14280 | 0.05626 | 80 | 2.538 | 0.0131 |
|  | consec_Other | 0.03708 | 0.01066 | 80 | 3.479 | 0.0008 |
|  | (consec_Other)^2 | -0.00086 | 0.00056 | 80 | -1.527 | 0.131 |
| RPR | Intercept | 0.24484 | 0.07922 | 75 | 3.091 | 0.0028 |
|  | consec_Other | 0.04413 | 0.01320 | 75 | 3.343 | 0.0013 |
|  | (consec_Other)^2 | -0.00068 | 0.00054 | 75 | -1.262 | 0.211 |

**Table 2:** Generalized Linear Mixed-Effects Model (GLMM, negative binomial) of poke counts across paradigms.

| Effect | Estimate | Std. Error | z value | p-value |
| --- | --- | --- | --- | --- |
| (Intercept, FR) | 6.370 | 0.056 | 113.81 | <2e-16 *** |
| Paradigm 2x2 | 0.653 | 0.078 | 8.36 | <2e-16 *** |
| Paradigm 5x5 | 0.664 | 0.078 | 8.50 | <2e-16 *** |
| Paradigm RPR | 1.635 | 0.077 | 21.12 | <2e-16 *** |

**Table 3:** Generalized Linear Mixed-Effects Model (GLMM, negative binomial) of other-poke counts across paradigms.

| Effect | Estimate | Std. Error | z value | p-value |
| --- | --- | --- | --- | --- |
| (Intercept, FR) | 3.332 | 0.114 | 29.25 | <2e-16 *** |
| Paradigm 2x2 | 3.100 | 0.138 | 22.41 | <2e-16 *** |
| Paradigm 5x5 | 3.117 | 0.138 | 22.53 | <2e-16 *** |
| Paradigm RPR | 4.477 | 0.138 | 32.57 | <2e-16 *** |

**Table 4:** Generalized Linear Mixed-Effects Model (GLMM, negative binomial) of pellet counts across paradigms.

| Effect | Estimate | Std. Error | z value | p-value |
| --- | --- | --- | --- | --- |
| (Intercept, FR) | 6.319 | 0.045 | 140.18 | <2e-16 *** |
| Paradigm 2x2 | -0.106 | 0.064 | -1.65 | 0.098 . |
| Paradigm 5x5 | -0.102 | 0.064 | -1.59 | 0.113 |
| Paradigm RPR | -0.047 | 0.064 | -0.74 | 0.460 |

### Effectiveness of Effort in Pellet Retrieval

To quantify how effectively mice converted their poking effort into rewards, we analyzed the instantaneous frequency of pellet retrieval across the different paradigms (Supplementary Figure 2). Four panels, arranged from top to bottom corresponding to the FR, 2×2, 5×5, and RPR paradigms, show the instantaneous frequency of pellet retrieval for the randomly selected mice in Figure 1. The instantaneous frequency for each pellet was calculated as the reciprocal of the time interval between the current pellet and the previous pellet. Across all paradigms, the instantaneous frequency of pellet retrieval was markedly lower and more stable than the instantaneous frequency of poking behavior. Unlike poking, which exhibited substantial variability and paradigm-dependent structure, pellet retrieval occurred at a relatively constrained rate with limited fluctuation over time. This pattern was consistently observed in the FR, 2×2, 5×5, and RPR paradigms, indicating that despite large differences in response dynamics, the temporal structure of reinforcement remained comparatively uniform across task designs. This stability in reward rate despite varying effort demands echoes findings from studies of behavioral regulation and homeostasis (McFarland and Sibly 1975, Boulos and Terman 1980).

### Aggregate Comparison Across All Mice and Paradigms

To assess the cumulative impact of response cost on effort and reward acquisition, we compared total pokes, other (incorrect) pokes, and pellet retrieval counts across all schedules (Figure 3A; n = 5 mice per paradigm). Total poke counts increased monotonically with task difficulty, with the FR schedule yielding the lowest counts and the RPR schedule the highest; the 2×2 and 5×5 schedules showed intermediate and comparable levels. A parallel pattern was observed for other pokes, indicating that unrewarded effort scales with schedule demands. In contrast, pellet retrieval counts remained stable across all four paradigms, suggesting that despite escalating costs, mice maintained consistent food intake.

**Figure 3:**
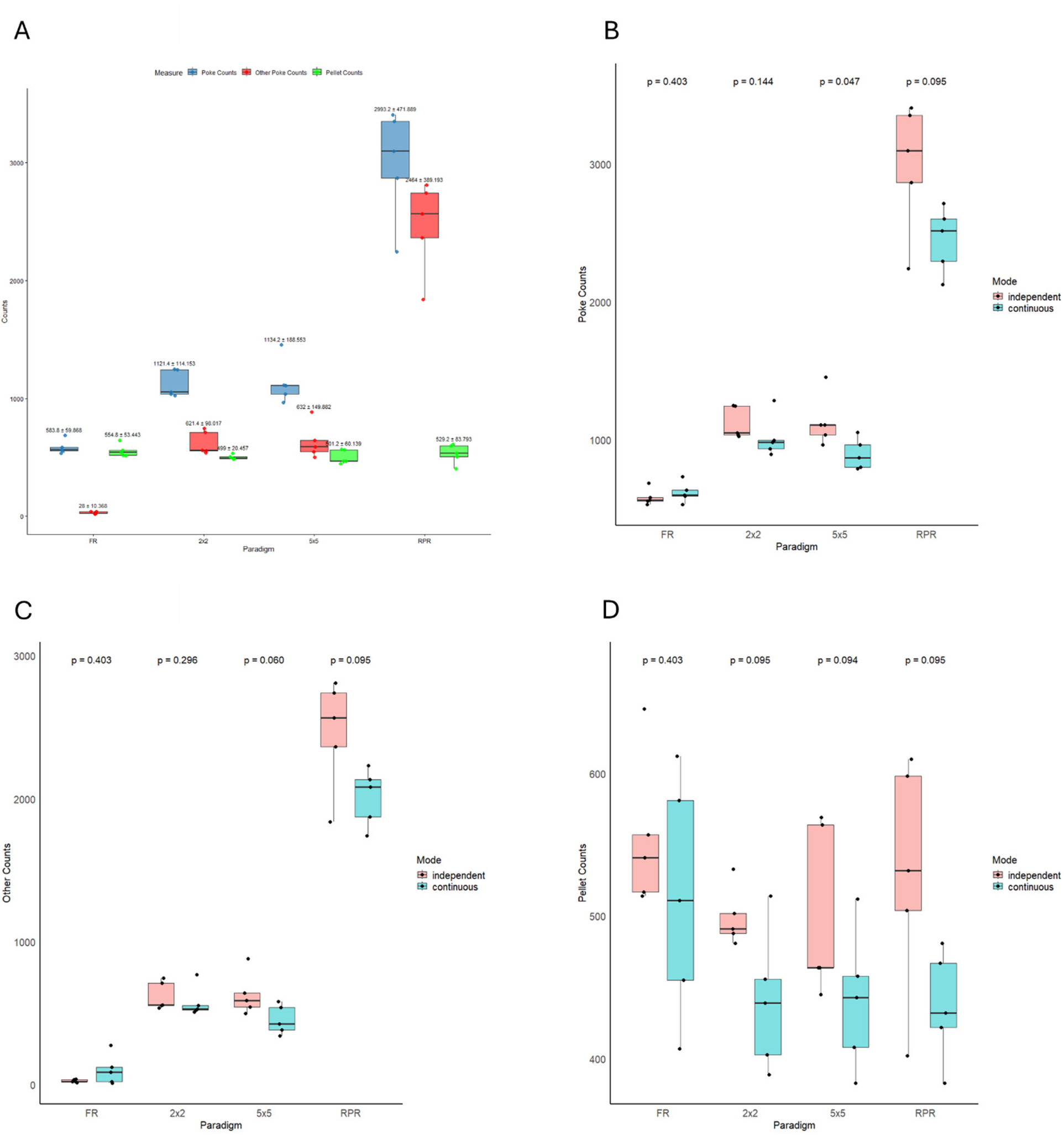
Poke and pellet counts across paradigms and training types. **(A) Total poke, other poke, and pellet retrieval counts for independent training.** Data are shown for all four paradigms, with five mice per paradigm. For each paradigm, three color-coded boxplots represent total poke count, other poke count, and pellet retrieval count. Each box summarizes data from five mice trained independently in that paradigm. **(B–D) Comparison of independent versus continuous training.** Each panel shows one type of count: **(B) total pokes**, **(C) other pokes**, and **(D) pellet retrievals**. Within each paradigm, two boxplots are shown: one for mice trained independently and one for mice trained continuously. Each box summarizes data from five mice. Statistical comparison between independent and continuous training within each paradigm was performed using Mann–Whitney tests, with p-values shown above each pair of boxes.

Generalized linear mixed-effects models (GLMM) with a negative binomial distribution confirmed these observations. Relative to the FR baseline, total poke counts were significantly elevated in all schedules (2×2: estimate = 0.653, p < 2e-16; 5×5: estimate = 0.664, p < 2e-16; RPR: estimate = 1.635, p < 2e-16). Similarly, other-poke counts were markedly higher in all schedules compared to FR (2×2: estimate = 3.100, p < 2e-16; 5×5: estimate = 3.117, p < 2e-16; RPR: estimate = 4.477, p < 2e-16). Crucially, pellet retrieval counts did not differ significantly from FR in any schedule (2×2: p = 0.098; 5×5: p = 0.113; RPR: p = 0.460), confirming that reward acquisition rate was preserved despite increased effort and failure tolerance. Together, these results demonstrate that mice adapt to rising behavioral obstacles by investing greater unrewarded effort, yet successfully defend baseline levels of goal attainment.

### Training History Does Not Confound Behavioral Phenotypes

To rule out order effects, we compared behavioral output between mice trained on independent versus continuous schedules. Mann–Whitney tests revealed no significant differences in total pokes, other pokes, or pellet counts within any paradigm (Figures 3B–3D; all p > 0.05), confirming that task history did not influence performance and that subsequent analyses could be collapsed across training cohorts.

### The PERCS Framework Reveals Multidimensional Effort Strategies

We next applied the PERCS framework to quantify behavior across five core dimensions—Persistence (P), Endurance (E), Resistance to Extinction (R), Consistency (C), and Sequence Stability (S)—with Resistance to Extinction grounded in behavioral momentum theory via devaluation-based satiety probes (Nevin and Grace 2005, Podlesnik and Shahan 2008). In the FR schedule, persistence bouts were virtually absent, yielding near-zero values for P, E, R, and C, while S approached ceiling (near 100% correct pokes), reflecting a habitual, low-effort response structure(Squire and Zola 1996, Graybiel 2008). The 2×2 and 5×5 schedules displayed similar multidimensional profiles, characterized by elevated P, E, and R alongside reduced S, indicating that alternating response demands require sustained effort and strategic endurance at the expense of sequence stability. The RPR schedule exhibited the lowest S—consistent with the mathematical unpredictability of a random environment that precludes stable sequence learning—while showing the highest P and R, reflecting maximal effort and continued responding despite uncertainty. Notably, RPR also revealed marked inter-individual variability in E and C, suggesting that high-demand conditions unmask divergent strategies in temporal consistency and endurance.

### PERCS Profiles Are Robust Across Training Histories

To assess reproducibility, we examined PERCS profiles separately in the continuous training cohort (Figure 4B) and in the combined dataset (Figure 4C). In both cases, the multidimensional signatures remained qualitatively identical to those observed in the independent cohort: FR was dominated by high S and minimal persistence-related dimensions, 2×2 and 5×5 showed intermediate elevation of P, E, and R with reduced S, and RPR exhibited low S with maximal P and R. The stability of these profiles across independent and continuous training histories demonstrates that PERCS dimensions capture stable behavioral phenotypes shaped by current task contingencies rather than by training artifacts.

**Figure 4:**
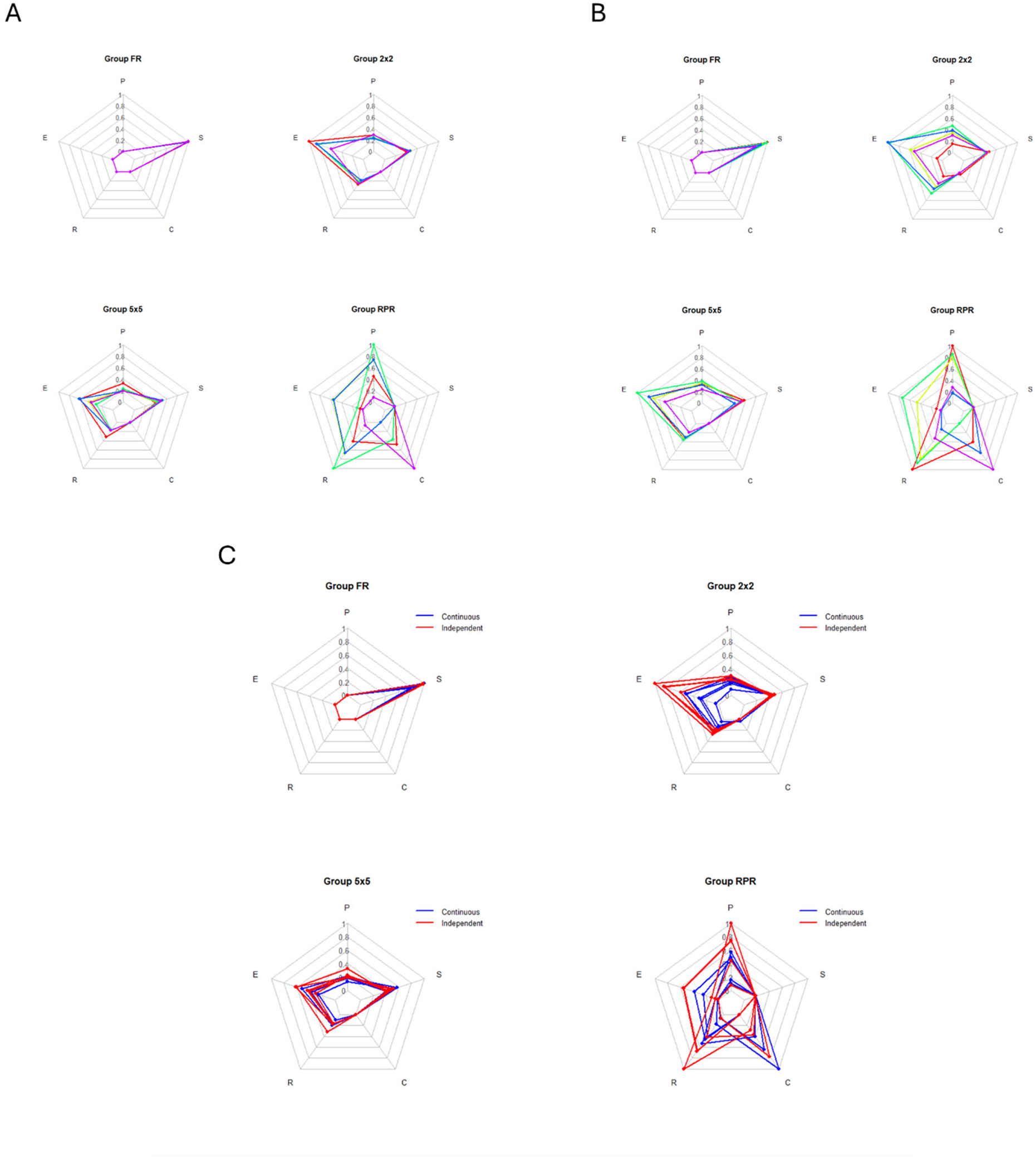
Radar plots of PERCS metrics across paradigms and training types. (A) Independent training paradigms. Radar plots illustrate PERCS profiles derived from mouse behavioral data obtained under independent paradigm training. The dataset includes 20 mice in total, with five mice per paradigm. Colors represent individual mice within each paradigm. For each PERCS dimension, values were normalized such that the maximum value across mice was set to 1, and all other values were scaled relative to this maximum. All dimensions are therefore constrained to the range 0–1, allowing direct comparison across dimensions and paradigms. **(B) Continuous training paradigms**. Radar plots show PERCS profiles for mice trained under continuous paradigms. Normalization and plotting conventions are the same as in panel A. **(C) Comparison of independent and continuous training.** Radar plots display PERCS profiles for mice across both independent and continuous training paradigms. Normalization and plotting conventions are identical to panels A and B, enabling direct comparison across training types.

### Multivariate Analysis Reveals Paradigm-Specific Persistence Architectures

To determine whether distinct task contingencies recruit dissociable combinations of persistence dimensions, we performed multivariate analyses of PERCS profiles. Analysis of multivariate dispersion revealed significant differences in within-paradigm variability across schedules (global permutation test, 𝑝 < 0.05; Table 5). RPR exhibited significantly greater dispersion than FR (𝑝 < 0.05), while comparisons between RPR and the alternating schedules (2×2, 𝑝 = 0.119; 5×5, 𝑝 = 0.151) did not reach statistical significance, likely reflecting limited sample size (n = 5 per paradigm) for high-dimensional comparisons. While this lack of significance with the alternating schedules is acknowledged as a limitation of the current sample size, the consistent pattern of median deviations (Figure 5A) suggests that RPR may drive greater individual variability in persistence strategies. This trend is conceptually significant, as it implies that environments with high uncertainty and effort demands may unmask or amplify pre-existing individual differences in coping strategies, rather than producing a uniform behavioral response. Nonetheless, median deviations (Figure 5A) indicated that RPR consistently showed the greatest overall spread, suggesting that high-demand, stochastic schedules amplify individual variability in persistence strategies.

**Figure 5:**
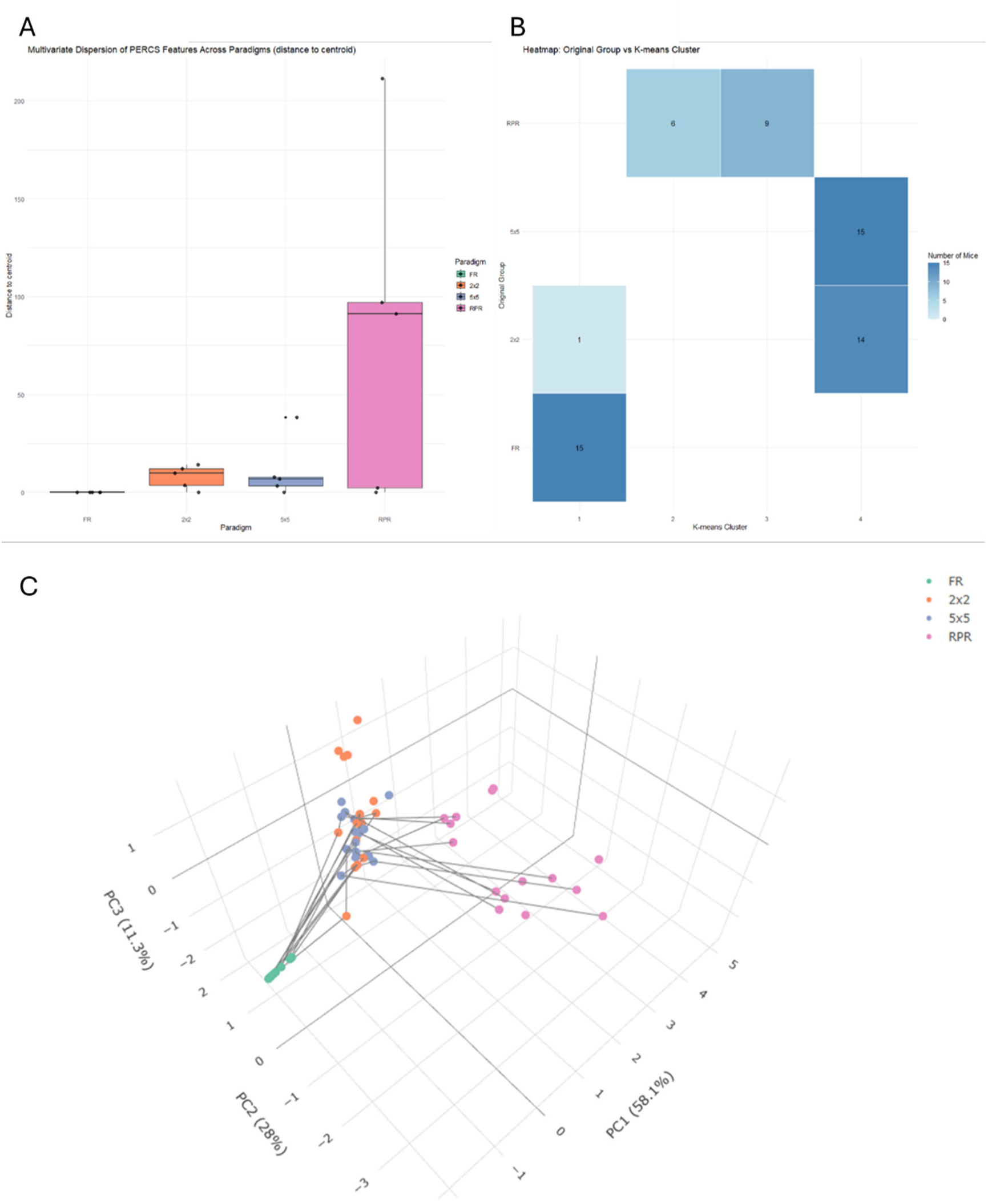
Structure and variability of mouse PERCS behavioral features across paradigms. (A) PERCS feature dispersion. Each point represents the Euclidean distance between an individual mouse’s 5-dimensional PERCS vector (P, E, R, C, S) and the centroid of its paradigm (FR, 2×2, 5×5, RPR). Boxes summarize the distribution of distances within each group. Group differences were assessed using a permutation-based test for dispersion (Anderson, 2006).**(B) Heatmap of k-means clustering.** The y-axis shows the original paradigms (FR, 2×2, 5×5, RPR), and the x-axis shows the k-means cluster assignment (1–4) of each mouse based on their 5-dimensional PERCS features. Each square represents the number of mice in that combination of original paradigm and assigned cluster, with color intensity indicating the count. **(C) Principal component analysis (PCA) of PERCS features.** Axes represent the first three principal components (PC1, PC2, PC3) of the 5-dimensional PERCS features across all mice. Points are colored by paradigm (FR, 2×2, 5×5, RPR). Lines connect repeated measurements from the same mouse in continuous training, illustrating within-mouse trajectories, whereas mice from independent training appear as single points.

**Table 5:** Global and pairwise permutation tests for multivariate dispersion of PERCS vectors across paradigms.

| Comparison | F | p-value | Interpretation |
| --- | --- | --- | --- |
| Overall (all 4 paradigms) | 3.56 | 0.001 | Significant difference in dispersion among groups |
| FR vs RPR | 4.29 | 0.001 | RPR significantly more dispersed than FR |
| 2x2 vs RPR | 3.47 | 0.119 | No significant difference |
| 5x5 vs RPR | 3.07 | 0.151 | No significant difference |

Unsupervised clustering of raw PERCS vectors (𝑘 = 4) yielded three empirically distinct groupings (Figure 5B). Cluster 1 comprised 16 mice, 94% of which (15 of 16) were from the FR schedule, confirming FR as a qualitatively distinct, low-effort phenotype. Clusters 2 and 3 consisted exclusively of RPR mice, revealing that the RPR schedule supports at least two separable behavioral subtypes. This divergence within the RPR schedule further supports the idea that high-demand conditions reveal distinct behavioral phenotypes, a finding that aligns with the trend towards increased multivariate dispersion observed in Figure 5A. Cluster 4 contained only 2×2 and 5×5 mice, indicating that these intermediate schedules occupy an overlapping region of PERCS space and recruit similar multidimensional persistence profiles.

Principal component analysis (PCA) corroborated this structure (Figure 5C). FR formed a compact, isolated cluster. The 2×2 and 5×5 schedules extensively overlapped, forming a second cluster. RPR displayed the widest dispersion, occupying a region partially separated from the other paradigms, consistent with its elevated multivariate variability. Together, these multivariate patterns demonstrate that different task contingencies evoke distinct combinations of persistence dimensions, supporting theoretical frameworks that posit multiple, dissociable mechanisms underlying persistent behavior (Carver and Scheier 1982, Eisenberger 1992)..

## Discussion

Our findings demonstrate that operant conditioning schedules delivered via the FED3 system produce a systematic, quantifiable continuum of persistent food-seeking behaviors. This continuum is directly shaped by the interplay of effort cost, sequential rule complexity, and outcome uncertainty inherent in each schedule, validating the hypothesis that persistence is not a binary state but a multidimensional behavioral phenotype that can be experimentally sculpted. The results align with and extend prior work using operant paradigms to study persistence. The finding that higher effort costs, as in the Random Progressive Ratio (RPR), elevate overall response output (Perseverance of Effort, P) and the frequency of persistence bouts is consistent with progressive ratio (PR) breakpoint assays, a gold standard for measuring motivational strength ((Hodos 1961, Richardson and Roberts 1996)). However, traditional PR schedules measure only the terminal point of effort expenditure. Our multidimensional analysis reveals that the RPR schedule not only increases P but also drastically reduces Repetitive Sequence Stability (S), a dimension not captured by a breakpoint. This aligns with behavioral momentum theory (Nevin and Grace 2000), which posits that behavior under rich, predictable reinforcement histories becomes more resistant to change. Our RPR schedule, by introducing high uncertainty, weakens this “behavioral mass,” leading to less structured, more variable responding, a profile captured by low S, a pattern that reflects the mathematical constraints of a random environment where stable sequence learning is impossible, thereby capturing effortful but unstructured responding. The structured alternation schedules (2×2, 5×5) produced a distinct profile, with elevated Strategic Endurance (E) and intermediate S. This mirrors findings from complex sequential learning and set-shifting tasks, where animals must maintain a rule in working memory and execute a specific action chain (Floresco, Zhang et al. 2009). We found a quadratic (inverted-U) relationship between consecutive unrewarded pokes and instantaneous frequency in simpler tasks (FR, 2×2), revealing an initial effort invigoration followed by strategic disengagement at very high failure counts. In contrast, high-demand tasks (5×5, RPR) maintained linear effort increases. The significant quadratic relationship between consecutive unrewarded pokes and instantaneous frequency in the 2×2 paradigm suggests an “inverted-U” of effort, an initial increase in vigor following failure, consistent with Amsel’s frustration theory (Amsel and Roussel 1952)), followed by a strategic disengagement at very high failure counts. This nuanced pattern would be invisible in a simple response-count analysis.

A key novel finding is that persistence bouts are triggered by unrewarded effort, not reward delivery. This was most evident in the RPR schedule, where high-rate poking continued even after occasional rewarded trials, and in the paired comparisons showing that “other” (incorrect) pokes had higher instantaneous frequencies than correct pokes in most paradigms. This directly challenges a purely reinforcement-driven account of persistence, which would predict that rewarded actions are the primary drivers of behavioral vigor. Instead, it strongly supports a motivational view where the absence of expected reward (frustrative nonreward) energizes ongoing behavior (Amsel and Roussel 1952). This mechanistic insight into the trigger of persistence is a significant advancement beyond descriptive schedules like PR, positioning frustrative nonreward as a primary, rather than secondary, engine of persistent responding. This energizing effect is most directly captured by the Perseverance of Effort (P) dimension, which quantifies total work output, and the inverted-U pattern in simpler tasks reveals the dynamic interplay between P and Strategic Endurance (E), where initial invigoration gives way to adaptive disengagement. This directly challenges a purely reinforcement-driven account of persistence and supports a motivational view where the absence of expected reward (frustrative nonreward) energizes ongoing behavior (Amsel and Roussel 1952). This mechanistic insight into the trigger of persistence is a significant advancement beyond descriptive schedules like PR. Furthermore, our use of Gaussian mixture modeling (GMM) to define persistence periods session-by-session provides objective, data-driven alternative to arbitrary frequency thresholds, enhancing the reproducibility of “persistence” as a behavioral construct. This method accounts for individual and session variability, making the identification of “persistent states” more robust and comparable across subjects and time, a methodological improvement over fixed criteria.

Finally, the Fixed Ratio (FR) schedule served as a critical control, generating a PERCS profile of near-zero P, E, R, and C, but maximal S. This profile represents a pure, efficient habit (Graybiel 2008), highly automated, temporally irregular, and requiring minimal cognitive engagement. It starkly illustrates that high sequence stability can exist independently of other persistence dimensions, a dissociation made explicit by our framework. The neural substrates of such habitual behavior likely involve corticostriatal loops and dopamine signaling (Jin, Tecuapetla et al. 2014, Goedhoop, van den Boom et al. 2022, Goedhoop, Arbab et al. 2023). In summary, our FED3 results confirm that schedule parameters dissociably shape distinct facets of persistence. They bridge classic operant concepts (breakpoint, resistance to extinction) with a richer, multidimensional characterization, providing a more precise map of how challenge structure dictates behavioral strategy.

Understanding behavioral persistence is of paramount importance. It is a transdiagnostic feature spanning adaptive success, from foraging and migration to academic and professional achievement, and debilitating psychopathology, including the compulsions of OCD, the apathy of depression (Marin 1991, Treadway and Zald 2011), the perseveration of frontal lobe syndromes (Sandson and Albert 1984), the inattention of ADHD (Barkley 1997), and the cognitive rigidity of ASD ((Graybiel and Rauch 2000); (Treadway and Zald 2011, Lord, Brugha et al. 2020)). Despite this centrality, the field has been historically fragmented, as detailed in our introduction. This fragmentation arises from divergent disciplinary roots: behaviorism’s focus on reinforcement history (Nevin and Grace 2000), human psychology’s emphasis on traits like grit (Duckworth, Peterson et al. 2007) and willpower (Baumeister, Bratslavsky et al. 1998), and clinical neuroscience’s categorical approach to disorders. Consequently, the field lacks a common operational language. The “persistence” measured by a PR breakpoint in a rat, the “tenacity” of a student studying for exams, and the “perseveration” of a patient with frontal lobe damage are often discussed in isolation, impeding the integration of knowledge across levels of analysis and the translation of mechanistic insights from model systems to human conditions.

Our work directly addresses this gap. By employing the FED3, a tool capable of inducing a full continuum of persistent states under controlled conditions, we move persistence from a descriptive phenomenon observed in response to external manipulations (e.g., threat, drugs) to an experimentally tractable dependent variable. This allows for the systematic investigation of its causes and components, a necessary step for a mature science.

The PERCS framework represents more than a novel description of our data; it offers a transformative roadmap for future research in behavioral neuroscience, ethology, and psychiatry. First, it provides a dimensional, quantitative phenotyping tool. Notably, our framework identifies Persistence (P), Endurance (E), Resistance to Extinction (R), Consistency (C), and Sequence Stability (S) as core dimensions, with Resistance to Extinction (R) measured through devaluation-based satiety probes, grounding this dimension in behavioral momentum (Nevin and Grace 2005, Podlesnik and Shahan 2008). PERCS offers a common metric for cross-species, cross-paradigm, and cross-disorder comparison. Future studies can now quantify whether a genetic mutation, pharmacological treatment, or environmental insult specifically alters Perseverance of Effort (P) versus Repetitive Sequence Stability (S). For example, does a dopamine agonist increase effort tolerance (P) in a PR task and strategic vigilance (E) in a sustained attention task? Such questions can now be addressed using mouse models of neuropsychiatric disorders(Peca, Feliciano et al. 2011, Carcea, Caraballo et al. 2021). PERCS allows these ostensibly different outcomes to be recognized as potentially related manipulations of the persistence spectrum. This dimensional approach aligns with the NIH’s Research Domain Criteria (RDoC) initiative (Onnela and Rauch 2016, Insel 2018),moving beyond diagnostic labels toward dysfunction in core behavioral systems. Second, PERCS enables disentangling adaptive from maladaptive persistence. A core promise of PERCS is its ability to distinguish functional persistence from dysfunctional perseveration. As shown in our results and theoretical model, pathology often involves an imbalance or extreme expression of one dimension (e.g., pathologically high R and S in OCD) or a deficit in another (e.g., pathologically low P and E in depression). By profiling patients along these five dimensions, clinicians could move toward a more precise, mechanism-based diagnosis. Treatment could then be targeted: cognitive-behavioral therapy with exposure and response prevention might specifically aim to reduce maladaptive R and S in OCD(Azrin and Nunn 1973, Davis, Ressler et al. 2006) (Azrin and Nunn 1973; Davis, Ressler et al. 2006), while behavioral activation therapy aims to augment deficient P and E in depression(Lejuez, Hopko et al. 2011).Cognitive remediation approaches (Wykes, Huddy et al. 2011) and executive function training could target specific dimensional deficits. Third, PERCS facilitates bridging levels of analysis from behavior to circuit. The dissociable nature of PERCS dimensions suggests they may have partially distinct neural substrates. P (effort) is heavily linked to dopamine signaling in the ventral striatum and anterior cingulate cortex (Salamone et al., 2007). E (endurance/attention) relies on norepinephrine and prefrontal networks (Aston-Jones and Cohen 2005). S (sequence stability) involves corticostriatal loops responsible for habit formation (Jin, Tecuapetla et al. 2014). C (temporal consistency) engages circadian and ultradian rhythm generators (Boulos and Terman 1980, Frank, Kupfer et al. 2005, Wang and Sun 2024). Future research can test these mappings directly by correlating neural activity (e.g., via calcium imaging or electrophysiology in mice performing our FED3 schedules) with simultaneously derived PERCS profiles. This would allow us to move from asking “what brain region is active during a hard task?” to “what circuit dynamic encodes a specific dimension of persistent challenge?” Fourth, PERCS enables computational psychiatry and personalized prediction. The tiered normalization strategy of PERCS allows raw behavioral data from diverse sources, home-cage monitoring, clinical task batteries, even digital phenotyping from smartphones (Barnett and Onnela 2020), to be transformed into comparable dimension scores. This opens the door for large-scale computational phenotyping. Machine learning models could be trained on these multidimensional profiles to predict individual risk for disorders, trajectory of illness, or response to specific treatments, fostering a new era of personalized medicine. While the current study establishes the normative adaptive landscape across these schedules, a critical next step involves identifying the “pathological tipping point” where adaptive persistence transitions into maladaptive perseveration. Although diseased models were initially considered, future research will utilize the baseline established here to explicitly profile such models along the PERCS dimensions to differentiate, for example, high S in OCD from low P in depression. While the current study establishes the normative adaptive landscape across these schedules, a critical next step involves identifying the “pathological tipping point” where adaptive persistence transitions into maladaptive perseveration. The diseased model mice mentioned in the Methods were part of a preliminary exploration, but the data are not included here as the focus of this foundational paper is to establish the normative PERCS framework and validate its sensitivity to schedule parameters. Future research will explicitly profile such disease models along the PERCS dimensions using the baseline established here to differentiate, for example, high S in OCD from low P in depression.

A limitation of the current study is the relatively small cohort used for independent training (N=5 per paradigm). While the primary effects on total pokes and dimension rankings were highly significant, the smaller sample size likely reduced the power of our multivariate dispersion tests. This is particularly relevant for the comparisons between RPR and the alternating schedules (2×2 and 5×5) in Figure 5A, where non-significant p-values may reflect a Type II error rather than a true absence of effect. The consistent pattern of median deviations and the emergence of two distinct RPR clusters in Figure 5B, however, suggest that these trends are biologically meaningful and warrant investigation in larger cohorts. Environments with high uncertainty and effort demands may unmask pre-existing individual differences in coping strategies, leading to divergent behavioral phenotypes. Crucially, however, the multidimensional behavioral signatures we observed were robust across both independent and continuous training histories (Figures 3B-D, 4A-C), demonstrating that the PERCS profiles are stable phenotypes shaped by current task contingencies rather than mere training artifacts. This robustness strengthens confidence that our findings reflect fundamental properties of the schedules themselves. Future studies with larger populations will be required to further refine the statistical boundaries of these behavioral clusters. In conclusion, this study establishes a synergistic methodology: using programmable operant schedules to induce a persistence continuum and applying the multidimensional PERCS framework to characterize it. We have demonstrated that schedule uncertainty and structure dissociably shape core dimensions of persistent behavior, with unrewarded effort acting as a key trigger, challenging reinforcement-centric models and positioning frustrative nonreward as a primary engine of persistence. By providing a unified, quantitative, and neurobiologically-grounded model, the PERCS framework transcends historical fragmentation. It offers a powerful new lens through which to view goal-directed behavior, from the rhythmic lick of a mouse to the sustained striving of a human, and a precise toolkit for deconstructing its breakdown in disease. This work lays the foundation for a unified, mechanistic science of behavioral persistence.

## Methods and Materials

### Subject details

All experimental procedures were approved by the Institutional Animal Care and Use Committee (IACUC) and the Biosafety Committee of the University of Wyoming. 10 male and 10 female immunocompetent mice of postnatal day 101.5±23.3 days on a B6.Cg-Snap25tm3.1Hze/J background were included in the datasets. All mice were bred on a C57/BL6J background. Mice older than 30 days were housed with same-sex littermates or alone in a vivarium at 21-23 °C with 25%-30% humidity and a 12-h light/dark cycle.

### Food Restriction Procedure

To elicit goal-oriented behavior, we implemented a food restriction protocol not exceeding 30 days, with body weight maintained no lower than 85% of baseline, consistent with IACUC guidelines and established standards for behavioral neuroscience research (National Research Council 2003). Mice typically consume 10-15% of body weight daily (3-5 grams of food). Mice were acclimated over a 7-day period with gradual reduction to the target level of 2 grams per day (Chevee, Kim et al. 2023). To ensure adequate caloric reserves and mitigate nutritional deficits, infant formula soymilk (SMA Wysoy) was provided as a supplement during handling, post-behavioral training, and post-experimentation. During training and experimental sessions, food pellets served as positive reinforcement and contributed to the daily total allowance of 2 grams (Goltstein, Reinert et al. 2018). Mice were monitored daily throughout the restriction period, with body weight recorded pre- and post-consumption. Health assessments included systematic evaluation of skin coat condition, posture, urine and feces output, and activity level. Any mouse exhibiting signs of distress or body weight drop below 85% of baseline was immediately removed from the study and allowed to recover prior to any potential re-entry (National Research Council 2003).

### Behavioral apparatus

We used a FED3-based behavioral apparatus(Matikainen-Ankney, Earnest et al. 2021) to implement multiple feeding paradigms. The FED3 chamber consisted of three ports: two lateral nose-poke ports (left and right) and a central pellet port for food delivery and retrieval. Task rules governing pellet delivery were fully programmable, allowing precise control over the response-outcome contingencies.

### Operant feeding paradigms

In this study, four distinct paradigms were employed. The fixed-ratio (FR) control condition was a paradigm with minimal response cost, in which a pellet was delivered to the central port following a single nose poke to either lateral port. The alternating 2×2 paradigm required alternating activation of the left and right nose-poke ports; when one lateral port was active, a single nose poke to that port immediately triggered pellet delivery, followed by pellet retrieval, and this poke-pellet-retrieval sequence constituted one response cycle. Two such response cycles were required before the active port switched to the opposite side, after which the same two-cycle sequence was repeated. The alternating 5×5 paradigm followed the same alternating structure as 2×2 but required five consecutive response cycles before switching to the opposite port. The random progressive ratio (RPR) condition imposed the highest response cost, requiring animals to perform a variable total number of nose pokes across both ports before pellet delivery; the required number of pokes was randomly drawn from a uniform distribution between 1 and 10 and was re-sampled after each pellet retrieval. These schedules were designed to systematically vary effort cost, sequential complexity, and reward uncertainty, building on classic operant conditioning principles(Skinner 1988).

### Response classification

To characterize operant responses, pokes were categorized based on their effectiveness in obtaining pellets. A poke that matched the current programmed rule and successfully triggered pellet delivery was classified as a correct poke (or rewarded poke). All other pokes were classified as other pokes (or incorrect pokes), including those directed to an inactive port and unnecessary additional pokes made after a correct poke but before pellet retrieval.

### Experimental design

We conducted two main experimental designs. In the independent paradigm training, each of the four paradigms was trained independently with five healthy, naive mice per paradigm, resulting in a total of 20 mice. In the consecutive paradigm training, three healthy naive mice and three diseased naive mice were each trained consecutively through all four paradigms, allowing within-subject comparison across paradigms; this cohort comprised a total of six mice.

### Definition and detection of persistence periods

For mice undergoing reversal training, a persistence period was defined as follows: it begins when a mouse’s poke instantaneous frequency exceeds a high threshold, continues while the poke repeats at least four times without falling below a lower bound, and ends when the instantaneous frequency drops below that lower bound. If the poke repeats fewer than four consecutive times at a high instantaneous frequency, it is not considered a persistence period. This explains why the FR paradigm, characterized by low-frequency, single pokes, does not contain formal persistence periods. This definition captures the essence of behavioral bouts as units of persistent action(Markowitz, Gillis et al. 2023) (Jin, Tecuapetla et al. 2014, Wiltschko, Johnson et al. 2015, Weinreb, Pearl et al. 2024).

### Threshold determination using Gaussian mixture models

To determine the high and low instantaneous frequency thresholds used to define persistence periods objectively, we fitted a two-component Gaussian mixture model (GMM) separately for each mouse and for each training session, using the flexmix R package. This session-specific, mouse-specific approach avoids pooling data across sessions or individuals and accounts for both inter-individual and within-animal variability in poking behavior. Let *yᵢ* denote the instantaneous frequency of the i th poke within a given mouse and session. We assumed that *yᵢ* arises from a mixture of two normal distributions, *yᵢ ∼ π₁N(μ₁, σ₁²) + π₂N(μ₂, σ₂²)*, where *π₁* and π₂ = 1-π₁ are the mixture proportions, and *μₖ* and *σₖ* denote the mean and variance of component k, for k = 1,2 . Model parameters (*πₖ, μₖ, σₖ²*) were estimated using the expectation–maximization (EM) algorithm. For each poke i, the posterior probability of belonging to component k was computed as *Pr(zᵢ = k | yᵢ) = (πₖ N(yᵢ | μₖ, σₖ²)) / (∑ⱼ πⱼ N(yᵢ | μⱼ, σⱼ²))*, where *zᵢ* is a latent component indicator. Each poke was assigned to the component with the higher posterior probability. The high instantaneous frequency threshold was defined as High bound = (max{yᵢ | zᵢ = low} + min{yᵢ | zᵢ = high}) / 2, and the low instantaneous frequency threshold was defined as *Low bound = μₗₒw*. These session-specific thresholds were then used to identify persistence periods as described above.

### The PERCS model: a formal, cross-species definition of persistence

We formally define behavioral persistence as the functional maintenance of goal-directed behavior over time despite challenges, moving beyond topographic description to a functional analysis of behavioral regulation(Nevin and Grace 2000). The PERCS model quantifies this as a vector in a five-dimensional adaptive strategy space, where the overall behavioral profile *P* is: *P = [P, E, R, C, S]*. Each dimension is defined as a theoretically dissociable component of persistence, with quantification methods drawn from established, often cross-species, experimental paradigms. *Perseverance of Effort (P)* is the total work output or energy expenditure directed toward a goal against increasing effort costs or diminishing returns, reflecting the intensity of engagement with a behavioral final common path (Salamone, Correa et al. 2007). It is quantified as *P = Σ(Goal-Directed Responses)* or *P = Breakpoint* on a progressive ratio (PR) schedule (Hodos 1961, Richardson and Roberts 1996); the PR breakpoint operationalizes the point at which the cost of effort outweighs the common currency value of the reward, providing a ratio-scale measure (Killeen, Posadas-Sanchez et al. 2009). *Strategic Endurance (E)* is the sustained, focused application of effort toward a goal under conditions of acute challenge, high cognitive load, or frustrative nonreward (e.g., delay discounting, distraction). This measures the stability of the goal representation in working memory against competing valuations and is quantified as *E = f(Time-on-Task, Accuracy | High Difficulty or Distraction)*, often indexed by latency to succumb to distraction or quit a demanding task (Mischel, Shoda et al. 1989). *Resistance to Extinction (R)* is the tendency to continue emitting a previously reinforced response after the reinforcement contingency is withdrawn. This captures a fundamental learning dynamic where adaptive persistence must be distinguished from maladaptive failure to update behavior(Nevin and Grace 2000) and is quantified as *R = Number of Responses during Extinction* or as resistance to change via disruptor probes (e.g., pre-feeding, novel stimuli) within behavioral momentum theory. *Temporal Consistency (C)* is the rhythmic stability, regularity, and predictability of a goal-directed behavioral stream over extended time scales (days, weeks, seasons); high C indicates habit-like, automated patterning (Graybiel 2008) behavior (Boulos and Terman 1980, Frank, Kupfer et al. 2005, Wang and Sun 2024). It is quantified as C ≈ 1 / Variance(Inter-Response Intervals or Session-to-Session Frequency), where low temporal variance indicates high rhythmicity and schedule control. *Repetitive Sequence Stability (S)* is the structured, ordered chaining of behavioral modules (“syllables” or actions) into a stable, repeatable sequence. This dimension quantifies the syntax of action and its rigidity vs. flexibility (Markowitz, Gillis et al. 2023); (Wiltschko, Johnson et al. 2015) and is quantified as *S = Markov Chain Stability* (e.g., high transition probability from A→B→C) or low sequence entropy (Markowitz, Gillis et al. 2023); (Wiltschko, Johnson et al. 2015).

### PERCS framework quantification for the present study

For the current experiment, each PERCS dimension was operationalized using the collected behavioral data as follows. *Perseverance of Effort (P)* was quantified as the total number of persistence bouts, reflecting total effort investment across the session. *Strategic Endurance (E)* was defined as the proportion of time spent in persistence periods relative to the total duration of each training paradigm, multiplied by the proportion of correct pokes among all pokes; this measure captures the ability to sustain effective effort toward goal attainment. *Resistance to Extinction (R)* was operationalized as the number of persistence bouts occurring after satiety, defined here as the point at which the mouse had obtained 120 pellets, reflecting continued effort even after the primary goal was achieved. *Temporal Consistency (C)* was measured as the variance in the duration of individual persistence bouts, indexing the stability of persistence over time. *Repetitive Sequence Stability (S)* was quantified as the correct poke rate; under the task rules, higher correct poke rates necessarily reflect stronger and more stable sequential behavioral structure.

### Normalization: a tiered framework for cross-context and cross-species comparison

Raw PERCS metrics exist on disparate scales (counts, distances, entropy values). To enable unified quantitative comparison and create a common currency for persistence (Tinbergen, 1963), we introduce a tiered normalization framework. This transforms raw dimension scores into comparable, unitless indices ranging from 0 (minimal observed or plausible persistence) to 1 (maximal), relative to a defined reference frame. The core theoretical principle is that an organism’s persistence is represented as a normalized vector in five-dimensional space: *Pₙₒᵣₘ = [Pₙₒᵣₘ, Eₙₒᵣₘ, Rₙₒᵣₘ, Cₙₒᵣₘ, Sₙₒᵣₘ]*. Each score’s interpretation depends on the reference criterion used for normalization. Three normalization strategies are applied based on the scope of inference and research question. *Within-Study (Norm-Referenced) Normalization* is used for comparing experimental groups within a controlled cohort (e.g., mice on different reinforcement schedules). Raw scores are Z-scored or min-max scaled relative to the observed minimum and maximum across the study sample; this answers which treatment group shows relatively more or less of a persistence dimension (Cronbach and Meehl 1955). *Biologically Anchored (Criterion-Referenced) Normalization* is used for naturalistic data or when study samples may not capture the full biological range. Scores are scaled using a priori defined, species-typical, or theoretically plausible minima and maxima (e.g., physiological limits, observed species maxima). This prevents compression artifacts and asks how an individual’s persistence compares to the adaptive range for its phenotype. *Conceptual Strategy (Ideal-Type) Normalization* is used for cross-species or cross-context strategy comparison (e.g., mouse operant persistence vs. deer migration vs. human academic diligence). Here, normalized scores are mapped to a common conceptual scale (minimal–typical–maximal adaptive persistence) based on functional equivalence (Tinbergen 1972)). This constructs idealized persistence profiles, such as “efficient forager” or “compulsive responder”, to which real behavioral vectors can be compared. Implementation and validation require that the choice of method is a critical, theory-driven decision. A decision guide and complete algorithms are provided in Supplementary Methods S1. Validation follows psychometric principles, including sensitivity analysis and tests of convergent/discriminant validity across normalization tiers ((Cronbach and Meehl 1955). In the present study, for the independent training data (Figure 14), we applied within-study normalization. For each PERCS dimension, values were normalized such that the maximum value across all 20 mice was set to 1, and all other values were scaled relative to this maximum. All dimensions are thus constrained to the range of 0 to 1, facilitating direct comparison across dimensions and paradigms.

### Statistical analysis

For descriptive and comparative statistics, paired t -tests were used to compare the instantaneous frequency of correct vs. other pokes within each paradigm. To analyze the effect of consecutive unrewarded pokes on instantaneous frequency, we fitted a linear mixed-effects model separately for each paradigm: *Yᵢⱼ = β₀ + β₁Xᵢⱼ + β₂Xᵢⱼ² + bᵢ + ɛᵢⱼ*, where *Yᵢⱼ* = average instantaneous frequency for mouse i at a given consecutive-other-poke count, *Xᵢⱼ* = number of consecutive other pokes, *β₀, β₁, β₂* = fixed effects (intercept, linear, quadratic slope), *bᵢ* = random intercept for mouse i, and *ɛᵢⱼ* = random error. To compare total poke counts, other-poke counts, and pellet counts across paradigms and training designs, we fitted separate generalized linear mixed-effects models (GLMMs) with a negative binomial distribution to account for over-dispersed count data. Models included Paradigm and, where applicable, Treatment and their interaction as fixed effects, with mouse identity as a random intercept to account for repeated measures. Mann-Whitney U tests were used to compare outcomes (poke counts, other poke counts, pellet counts) between the independent and continuous training designs for each paradigm. All analyses were performed in R (version 4.x.x). A significance threshold of p<0.05 was used.

## Acknowledgements

We thank C. Zhang for animal husbandry, animal genotyping, histology assistance and items purchasing. This work is supported by grants from National Institute of Mental Health (1R21MH131363), and from National Institute of General Medical Sciences (2P20GM121310).

## Author contributions

Q.Q.S. designed the experiment and supervised the project, acquired the funding, and wrote the manuscript. T.C. analyzed data, performed statistical analysis and wrote the manuscript. W.J. S.C. and SC designed and performed FED3 experiments, analyzed the data and wrote the manuscript.

---

**Supplemental Figure 1:**
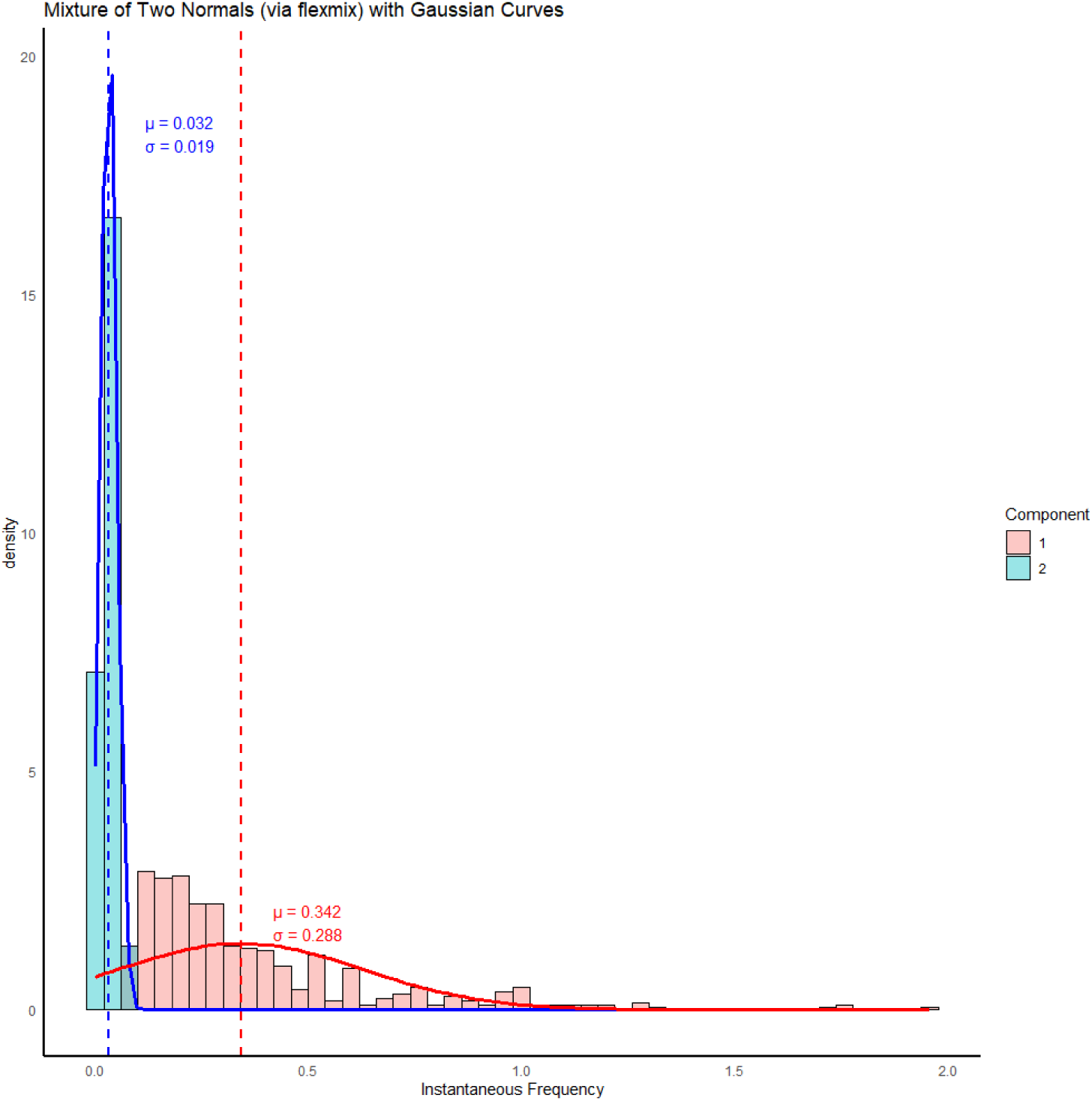
Example of a Gaussian mixture model for instantaneous poking frequency. The histogram shows the distribution of instantaneous poke frequency for a single mouse as an illustrative example. A Gaussian mixture model was applied to this distribution. The solid curves in blue and red represent the estimated normal components fitted to the histogram. The dashed vertical lines indicate the mean of each estimated normal distribution.

**Supplemental Figure 2:**
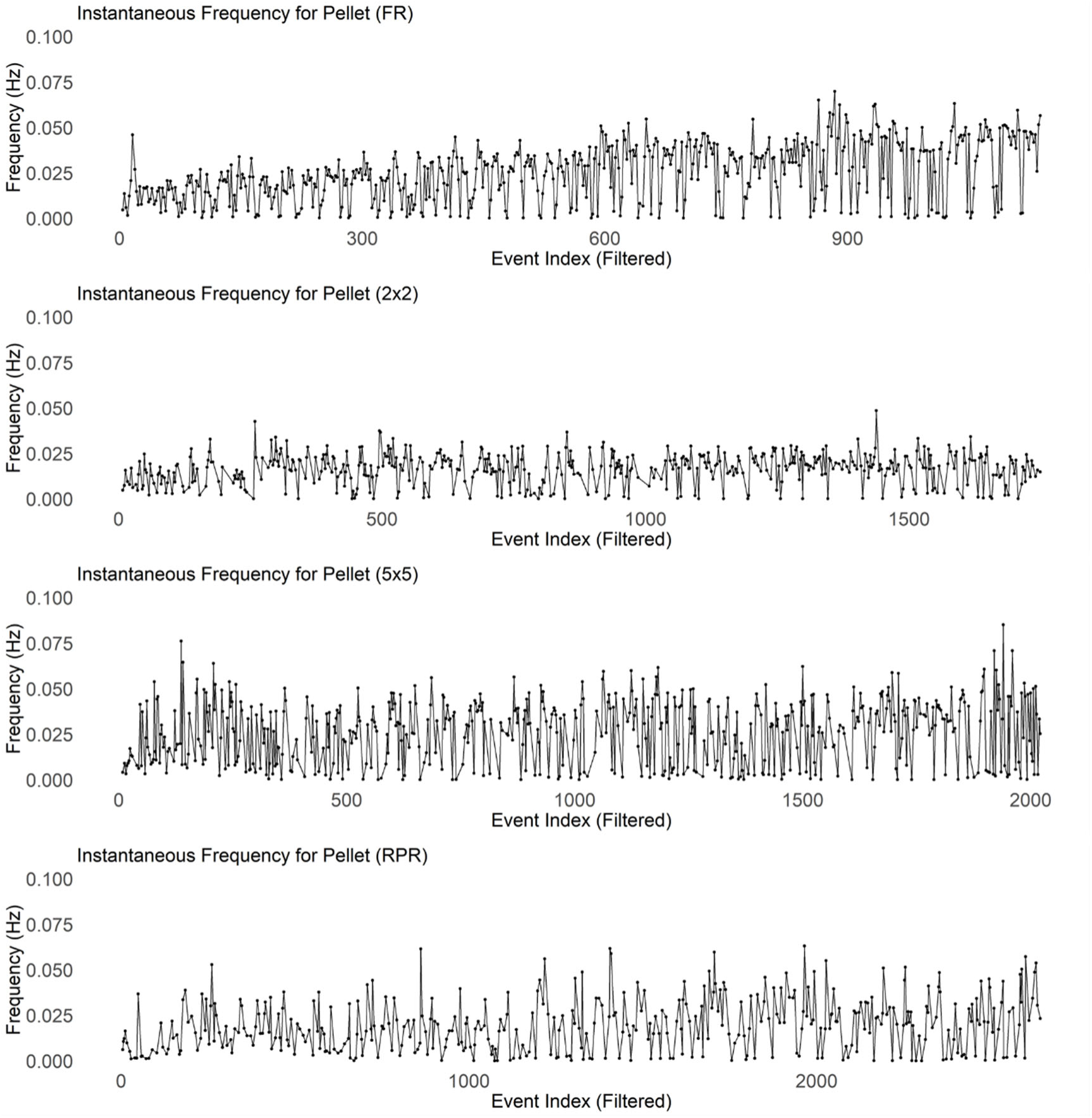
Instantaneous frequency of pellet retrieval. For each paradigm, the instantaneous pellet retrieval frequency was calculated as the reciprocal of the interval between consecutive pellet retrievals. Each dot represents a single pellet retrieval plotted over time. The panels are arranged from top to bottom corresponding to the four paradigms (FR, 2×2, 5×5, and RPR).

## References

1. Amsel, A. and J. Roussel (1952). “Motivational properties of frustration. I. Effect on a running response of the addition of frustration to the motivational complex.” J Exp Psychol 43(5): 363–366.

2. Aston-Jones, G. and J. D. Cohen (2005). “An integrative theory of locus coeruleus-norepinephrine function: adaptive gain and optimal performance.” Annu Rev Neurosci 28: 403–450.

3. Azrin, N. H. and R. G. Nunn (1973). “Habit-reversal: a method of eliminating nervous habits and tics.” Behav Res Ther 11(4): 619–628.

4. Barkley, R. A. (1997). “Attention-deficit/hyperactivity disorder, self-regulation, and time: toward a more comprehensive theory.” J Dev Behav Pediatr 18(4): 271–279.

5. Barnett, I. and J. P. Onnela (2020). “Inferring mobility measures from GPS traces with missing data.” Biostatistics 21(2): e98–e112.

6. Barrett, J. M., M. G. Raineri Tapies and G. M. G. Shepherd (2020). “Manual dexterity of mice during food-handling involves the thumb and a set of fast basic movements.” PLoS One 15(1): e0226774.

7. Baumeister, R. F., E. Bratslavsky, M. Muraven and D. M. Tice (1998). “Ego depletion: is the active self a limited resource?” J Pers Soc Psychol 74(5): 1252–1265.

8. Boulos, Z. and M. Terman (1980). “Food availability and daily biological rhythms.” Neurosci Biobehav Rev 4(2): 119–131.

9. Bradshaw, C. M. and P. R. Killeen (2012). “A theory of behaviour on progressive ratio schedules, with applications in behavioural pharmacology.” Psychopharmacology (Berl) 222(4): 549–564.

10. Carcea, I., N. L. Caraballo, B. J. Marlin, R. Ooyama, J. S. Riceberg, J. M. Mendoza Navarro, M. Opendak, V. E. Diaz, L. Schuster, M. I. Alvarado Torres, H. Lethin, D. Ramos, J. Minder, S. L. Mendoza, C. J. Bair-Marshall, G. H. Samadjopoulos, S. Hidema, A. Falkner, D. Lin, A. Mar, Y. Z. Wadghiri, K. Nishimori, T. Kikusui, K. Mogi, R. M. Sullivan and R. C. Froemke (2021). “Oxytocin neurons enable social transmission of maternal behaviour.” Nature 596(7873): 553–557.

11. Carver, C. S. and M. F. Scheier (1982). “Control theory: a useful conceptual framework for personality-social, clinical, and health psychology.” Psychol Bull 92(1): 111–135.

12. Carver, C. S. and M. F. Scheier (1998). On the self-regulation of behavior. New York, NY, US, Cambridge University Press.

13. Chevee, M., C. J. Kim, N. Crow, E. G. Follman, M. Z. Leonard and E. S. Calipari (2023). “Food Restriction Level and Reinforcement Schedule Differentially Influence Behavior during Acquisition and Devaluation Procedures in Mice.” eNeuro 10(9).

14. Cronbach, L. J. and P. E. Meehl (1955). “Construct validity in psychological tests.” Psychol Bull 52(4): 281–302.

15. Davis, M., K. Ressler, B. O. Rothbaum and R. Richardson (2006). “Effects of D-cycloserine on extinction: translation from preclinical to clinical work.” Biol Psychiatry 60(4): 369–375.

16. de Araujo Salgado, I., C. Li, C. J. Burnett, S. Rodriguez Gonzalez, J. J. Becker, A. Horvath, T. Earnest, A. V. Kravitz and M. J. Krashes (2023). “Toggling between food-seeking and self-preservation behaviors via hypothalamic response networks.” Neuron 111(18): 2899–2917 e2896.

17. Duckworth, A. L., C. Peterson, M. D. Matthews and D. R. Kelly (2007). “Grit: perseverance and passion for long-term goals.” J Pers Soc Psychol 92(6): 1087–1101.

18. Eisenberger, R. (1992). “Learned industriousness.” Psychol Rev 99(2): 248–267.

19. Floresco, S. B., Y. Zhang and T. Enomoto (2009). “Neural circuits subserving behavioral flexibility and their relevance to schizophrenia.” Behav Brain Res 204(2): 396–409.

20. Frank, E., D. J. Kupfer, M. E. Thase, A. G. Mallinger, H. A. Swartz, A. M. Fagiolini, V. Grochocinski, P. Houck, J. Scott, W. Thompson and T. Monk (2005). “Two-year outcomes for interpersonal and social rhythm therapy in individuals with bipolar I disorder.” Arch Gen Psychiatry 62(9): 996–1004.

21. Geurts, H. M., B. Corbett and M. Solomon (2009). “The paradox of cognitive flexibility in autism.” Trends Cogn Sci 13(2): 74–82.

22. Goedhoop, J., T. Arbab and I. Willuhn (2023). “Anticipation of Appetitive Operant Action Induces Sustained Dopamine Release in the Nucleus Accumbens.” J Neurosci 43(21): 3922–3932.

23. Goedhoop, J. N., B. J. G. van den Boom, R. Robke, F. Veen, L. Fellinger, W. van Elzelingen, T. Arbab and I. Willuhn (2022). “Nucleus accumbens dopamine tracks aversive stimulus duration and prediction but not value or prediction error.” Elife 11.

24. Goltstein, P. M., S. Reinert, A. Glas, T. Bonhoeffer and M. Hubener (2018). “Food and water restriction lead to differential learning behaviors in a head-fixed two-choice visual discrimination task for mice.” PLoS One 13(9): e0204066.

25. Graybiel, A. M. (2008). “Habits, rituals, and the evaluative brain.” Annu Rev Neurosci 31: 359–387.

26. Graybiel, A. M. and S. L. Rauch (2000). “Toward a neurobiology of obsessive-compulsive disorder.” Neuron 28(2): 343–347.

27. Hodos, W. (1961). “Progressive ratio as a measure of reward strength.” Science 134(3483): 943–944.

28. Insel, T. R. (2018). “Digital phenotyping: a global tool for psychiatry.” World Psychiatry 17(3): 276–277.

29. Jin, X., F. Tecuapetla and R. M. Costa (2014). “Basal ganglia subcircuits distinctively encode the parsing and concatenation of action sequences.” Nat Neurosci 17(3): 423–430.

30. Killeen, P. R., D. Posadas-Sanchez, E. B. Johansen and E. A. Thrailkill (2009). “Progressive ratio schedules of reinforcement.” J Exp Psychol Anim Behav Process 35(1): 35–50.

31. Lejuez, C. W., D. R. Hopko, R. Acierno, S. B. Daughters and S. L. Pagoto (2011). “Ten year revision of the brief behavioral activation treatment for depression: revised treatment manual.” Behav Modif 35(2): 111–161.

32. Lord, C., T. S. Brugha, T. Charman, J. Cusack, G. Dumas, T. Frazier, E. J. H. Jones, R. M. Jones, A. Pickles, M. W. State, J. L. Taylor and J. Veenstra-VanderWeele (2020). “Autism spectrum disorder.” Nat Rev Dis Primers 6(1): 5.

33. Marin, R. S. (1991). “Apathy: a neuropsychiatric syndrome.” J Neuropsychiatry Clin Neurosci 3(3): 243–254.

34. Markowitz, J. E., W. F. Gillis, M. Jay, J. Wood, R. W. Harris, R. Cieszkowski, R. Scott, D. Brann, D. Koveal, T. Kula, C. Weinreb, M. A. M. Osman, S. R. Pinto, N. Uchida, S. W. Linderman, B. L. Sabatini and S. R. Datta (2023). “Spontaneous behaviour is structured by reinforcement without explicit reward.” Nature 614(7946): 108–117.

35. Matikainen-Ankney, B. A., T. Earnest, M. Ali, E. Casey, J. G. Wang, A. K. Sutton, A. A. Legaria, K. M. Barclay, L. B. Murdaugh, M. R. Norris, Y. H. Chang, K. P. Nguyen, E. Lin, A. Reichenbach, R. E. Clarke, R. Stark, S. M. Conway, F. Carvalho, R. Al-Hasani, J. G. McCall, M. C. Creed, V. Cazares, M. W. Buczynski, M. J. Krashes, Z. B. Andrews and A. V. Kravitz (2021). “An open-source device for measuring food intake and operant behavior in rodent home-cages.” Elife 10.

36. McFarland, D. J. and R. M. Sibly (1975). “The behavioural final common path.” Philos Trans R Soc Lond B Biol Sci 270(907): 265–293.

37. Mischel, W., Y. Shoda and M. I. Rodriguez (1989). “Delay of gratification in children.” Science 244(4907): 933–938.

38. Nevin, J. A. and R. C. Grace (2000). “Behavioral momentum and the law of effect.” Behav Brain Sci 23(1): 73–90; discussion 90-130.

39. Nevin, J. A. and R. C. Grace (2000). “Preference and resistance to change with constant-duration schedule components.” J Exp Anal Behav 74(1): 79–100.

40. Nevin, J. A. and R. C. Grace (2005). “Resistance to extinction in the steady state and in transition.” J Exp Psychol Anim Behav Process 31(2): 199–212.

41. Onnela, J. P. and S. L. Rauch (2016). “Harnessing Smartphone-Based Digital Phenotyping to Enhance Behavioral and Mental Health.” Neuropsychopharmacology 41(7): 1691–1696.

42. Peca, J., C. Feliciano, J. T. Ting, W. Wang, M. F. Wells, T. N. Venkatraman, C. D. Lascola, Z. Fu and G. Feng (2011). “Shank3 mutant mice display autistic-like behaviours and striatal dysfunction.” Nature 472(7344): 437–442.

43. Podlesnik, C. A. and T. A. Shahan (2008). “Response-reinforcer relations and resistance to change.” Behav Processes 77(1): 109–125.

44. Richardson, N. R. and D. C. Roberts (1996). “Progressive ratio schedules in drug self-administration studies in rats: a method to evaluate reinforcing efficacy.” J Neurosci Methods 66(1): 1–11.

45. Salamone, J. D., M. Correa, A. Farrar and S. M. Mingote (2007). “Effort-related functions of nucleus accumbens dopamine and associated forebrain circuits.” Psychopharmacology (Berl) 191(3): 461–482.

46. Sandson, J. and M. L. Albert (1984). “Varieties of perseveration.” Neuropsychologia 22(6): 715–732.

47. Sayar-Atasoy, N., Y. Yavuz, C. Laule, C. Dong, H. Kim, J. Rysted, K. Flippo, D. Davis, I. Aklan, B. Yilmaz, L. Tian and D. Atasoy (2024). “Opioidergic signaling contributes to food-mediated suppression of AgRP neurons.” Cell Rep 43(1): 113630.

48. Skinner, B. F. (1988). “The operant side of behavior therapy.” J Behav Ther Exp Psychiatry 19(3): 171–179.

49. Squire, L. R. and S. M. Zola (1996). “Structure and function of declarative and nondeclarative memory systems.” Proc Natl Acad Sci U S A 93(24): 13515–13522.

50. Thorndike, E. L. (1933). “A Proof of the Law of Effect.” Science 77(1989): 173–175.

51. Tinbergen, N. (1972). “Functional ethology and the human sciences.” Proc R Soc Lond B Biol Sci 182(1069): 385–410.

52. Treadway, M. T. and D. H. Zald (2011). “Reconsidering anhedonia in depression: lessons from translational neuroscience.” Neurosci Biobehav Rev 35(3): 537–555.

53. Wang, Y. and Q. Q. Sun (2024). “A prefrontal motor circuit initiates persistent movement.” bioRxiv.

54. Weinreb, C., J. E. Pearl, S. Lin, M. A. M. Osman, L. Zhang, S. Annapragada, E. Conlin, R. Hoffmann, S. Makowska, W. F. Gillis, M. Jay, S. Ye, A. Mathis, M. W. Mathis, T. Pereira, S. W. Linderman and S. R. Datta (2024). “Keypoint-MoSeq: parsing behavior by linking point tracking to pose dynamics.” Nat Methods 21(7): 1329–1339.

55. Wiltschko, A. B., M. J. Johnson, G. Iurilli, R. E. Peterson, J. M. Katon, S. L. Pashkovski, V. E. Abraira, R. P. Adams and S. R. Datta (2015). “Mapping Sub-Second Structure in Mouse Behavior.” Neuron 88(6): 1121–1135.

56. Wykes, T., V. Huddy, C. Cellard, S. R. McGurk and P. Czobor (2011). “A meta-analysis of cognitive remediation for schizophrenia: methodology and effect sizes.” Am J Psychiatry 168(5): 472–485.

